# Extracellular vesicle molecular signatures characterize metastatic dynamicity in ovarian cancer

**DOI:** 10.1101/2021.04.22.440951

**Authors:** Amber Gonda, Nanxia Zhao, Jay V. Shah, Jake N. Siebert, Srujanesh Gunda, Berk Inan, Mijung Kwon, Steven K. Libutti, Prabhas V. Moghe, Nicola L. Francis, Vidya Ganapathy

**Author notes:** **Correspondence:** Vidya Ganapathy, Assistant Research Professor, Nicola L. Francis, Research Associate.

## Abstract

Late-stage diagnosis of ovarian cancer drastically lowers 5-year survival rate from 90% to 30%. Early screening tools that use non-invasive sampling methods combined with high specificity and sensitivity can significantly increase survival. Emerging research employing blood-based screening tools have shown promise in non-invasive detection of cancer. Our findings in this study show the potential of a small extracellular vesicle (sEV)-derived signature as a non-invasive longitudinal screening tool in ovarian cancer. We identified a 7-gene panel in these sEVs that overlapped with an established tissue-derived metastatic ovarian carcinoma signature. We found the 7-gene panel to be differentially expressed with tumor development and metastatic spread. While there were quantifiable changes in genes from the 7-gene panel in plasma-derived sEVs from ovarian cancer patients, we were unable to establish a definitive signature due to low sample number. The most notable finding was a significant change in the ascites-derived sEV gene signature that overlapped with that of the plasma-derived sEV signature at varying stages of disease progression. Taken together our findings show that differential expression of metastatic genes derived from circulating sEVs present a minimally invasive screening tool for ovarian cancer detection and longitudinal monitoring of molecular changes associated with progression and metastatic spread.

## Introduction

Ovarian cancer, is the fifth most deadly cancer in females due to its diagnosis at advanced stages of the disease (1, 2). Statistics show a 5-year survival rate of 90% when detected early (2) and of 30% when diagnosed at later stages (1). Almost 80% of ovarian cancer diagnoses occur at advanced stages due to its non-specific symptoms (2, 3) and lack of tumor-specific screening tools (4). Current screening tools such as transvaginal ultrasound can assess volume- and morphology-based changes but are non-specific, leading to false-positive outcomes (4–6). The measure of tumor biomarkers such as CA-125 has met with little success, due to an overwhelmingly high false-positive rate (7–10). Dochez et al. found that in order to improve on the current screening platforms that suffer from low specificity in early stages, assessing the levels of HE4 with CA-125 improves screening efficiency (11). Brodsky et al. identified a 6-gene signature that differentiates metastatic and primary ovarian lesions (12). While these show potential for gene signatures to predict disease progression, staging and treatment outcomes, most of these are done using samples collected from invasive biopsy-derived specimens (13–17).

Liquid biopsy, a concept that originated in 1948 with the definition of circulating DNA free from cells in human blood (18), has bridged the gap by providing a means for disease diagnosis, prognosis, and therapy decisions in the clinic independent of invasive tissue biopsies. The concept since then has evolved to include ribonucleic acids, circulating tumor cells, extracellular vesicles (EVs), and tumor educated platelets. Liquid biopsies can minimize the need for invasive tissue sampling while enabling longitudinal monitoring during the course of the disease. Most recently, the US Food and Drug Administration (FDA) has started approval of liquid biopsies as companion diagnostics (19). Cell-free DNA-based tests such as Guardant360 CDx, for non-small cell lung cancer, Signatera™, for minimal residual disease detection, FoundationOne^®^ Liquid CDx, for pan-tumor screen are amongst of the recently approved liquid biopsy platforms (20, 21). One of the most important improvements liquid biopsies offer over traditional tissue biopsies, is the potential to monitor tumor changes longitudinally (22, 23).

Extracellular vesicles (EVs) in particular hold significant promise in the successful application of liquid biopsies to the clinic. Small extracellular vesicles (sEVs), or exosomes, ranging in size from 40-160nm, have been shown to be effective carriers of functional proteins and nucleic acids to other cells both in the local environment and to distant sites (24). The contents of these vesicles are protected from the degrading circulatory environment and contain multiple molecular markers that are specific to the primary tumor and its microenvironment (25). ExoDx™ Prostate IntelliScore or EPI, an sEV-based test, for prostate cancer (26), is the first FDA-approved exosome-based test that focuses on patient stratification for conducting biopsies.

The challenge liquid biopsy-based screening tools face is validated demonstration of earlier detection than conventional screening tools. This requires large sample sizes and longitudinal tracking of a small fraction of the population that will develop cancer over time (27). Hence, we tested for the first-time the potential of a sEV-based ovarian signature to longitudinally predict disease progression in a mouse model of ovarian cancer metastases (**Box 1**). We explored known ovarian cancer metastatic genes extracted from several different gene data sets (12, 28–32) to combine targets from disparate studies of tissue biopsies and evaluate the possibility of using them as biomarkers in a liquid biopsy. These genes (*AEBP1, ACTB, COL11A1, COL5A1, LOX, NECTIN4, POSTN, SNAI1, THBS1, TIMP3*) as outlined by Cheon, et. al. (28) have a common functional goal of altering the tumor microenvironment (TME) through collagen remodeling. Collagen remodeling is a key event in metastasis and correlates with poor prognosis in multiple cancers (33). Collagen remodeling in ovarian cancer is thought to not only contribute to peritoneal metastases and ascites formation (28) but also to platinum drug resistance (34). In this study, we demonstrate that extracellular vesicles could be used not only in the identification of ovarian tumors, but more importantly to detect molecular changes that occur as the tumor progresses and metastasizes. We found differential expression of the 10-gene panel both in plasma and ascites-derived sEVs collected from a mouse model of metastatic ovarian cancer. The expression level changes in these genes correlated with tumor presence and longitudinal tumor progression. We demonstrated a part of our 10-gene signature to correlate with tumor presence in comparing serum-derived extracellular vesicle gene signatures from tumor-bearing versus healthy patients. This study reports on the first such feasibility demonstrated to date of small extracellular vesicles as a liquid biopsy tool for longitudinal monitoring/screening for ovarian cancer progression.

### Box1: Author summary

**Why was this study done?**

- Ovarian cancer, the second most common gynecological cancer, has a low survival rate primarily due to lack of diagnostic tools to detect tumors at early localized stages.
- Liquid biopsy is emerging as a powerful tool for non-invasive monitoring of tumor progression, metastases prediction, and therapy response.
- Small extracellular vesicles (sEV) have particularly gained attention due to their prevalence in all body fluids, biological stability, and their enhanced ability to capture biological information from parental cells compared to other analytes used in liquid biopsy.
- This was designed as the first such investigation to probe whether a sEV-based diagnostic would elicit tumor specific signatures in ovarian cancers and also follow the longitudinal changes in tumor progression through a change in sEV gene signature.

**What did the researchers do and find?**

- We established a 10-gene signature involving genes associated with collagen remodeling by validating their correlation to ovarian cancer prognosis using the OvMark dataset. The hazard ratio determined the correlation of gene expression to overall disease-free survival and resulting prognosis.
- Subsequently, using a mouse model of human ovarian cancer peritoneal metastases, we established the fidelity of the 10-gene signature to disease progression as metastases progressed over a three-week period. We found seven of the 10 genes in the signature at quantifiable levels in plasma-derived sEVs.
- When correlated with a small cohort of patient samples, there was a correlation between three of the seven genes in predicting metastases compared to normal serum.
- Additionally, ascites-derived sEVs from the mouse peritoneal metastases model exhibited a quantifiable increase in 5 of the 10 genes in the signature, with a correlation between the signatures from ascites-derived sEVs and plasma-derived sEVs at week 3 of tumor progression.

**What do these findings mean?**

- Our findings indicate for the first time the possibility of a sEV-derived signature as a diagnostic tool for ovarian cancer metastases prediction.
- The findings were based on a small cohort of animal and human samples and future work will validate this in a larger cohort of samples.
- The findings were based on plasma/serum-derived sEV from a tumor-bearing subject and future work with isolation of a cancer-specific sEV population will provide us with a reproducible signature without non-specific interference from non-cancerous sEVs.

## Methods

### Extracellular vesicle isolation

Extracellular vesicles were isolated from plasma and ascites from mice, and serum from human patients. Plasma/serum was isolated and collected from mice and from human patients. ExoQuick (Systems Biosciences Inc, Mountain View, CA) was used to isolate extracellular vesicles according to manufacturer’s instructions. Briefly, ExoQuick was added to 400-500 μl of plasma/serum at a 250 μl:63 μl ratio (plasma/serum to reagent), mixed thoroughly and incubated for 30min at 4°C. After centrifugation at 3,000 x g for 10 minutes, the pellet was resuspended in 1x PBS (Gibco, Thermo Fisher Scientific, Waltham, MA).

Ascites collected from mice was centrifuged at 2000 x g for 30 minutes at room temperature to remove cells and debris. The clarified supernatant (500μl-1ml starting volume) was mixed with a volume of Total Exosome Isolation-cell culture (Thermo Fisher Scientific, Waltham, MA) equal to half of the supernatant volume and incubated overnight at 4°C. The samples were then centrifuged at 10,000 x g for 10 minutes at 4°C. Pellets were resuspended in 1x PBS.

### Extracellular vesicle characterization

Size and concentration measurements were performed using a Malvern Nanosight NS300 (Malvern Panalytical, UK). Isolated extracellular vesicles were run at a 1:2000 dilution in PBS. Machine settings were as follows: Camera level: 11-12, data collection: 5×15sec, flow rate: 20, analysis setting: 6-8.

### RNA isolation

RNA was isolated from extracellular vesicles using a modified Trizol protocol. Trizol (Thermo Fisher Scientific, Waltham, MA) was added to each sample such that the sample volume was 10% of the Trizol volume. Samples were lysed and incubated for 5 minutes in Trizol and then 1-bromo-3-chloropropane (BCP) (Molecular Research Center, Inc. Cincinnati, OH) was added to separate the RNA from the remaining material. The RNA-containing aqueous phase was incubated with isopropanol and RNA was pelleted by centrifugation. The pellet was subsequently suspended in 75% ethanol and incubated overnight at −20°C. The next day the RNA was pelleted and washed by an additional incubation with 75% ethanol. After the second centrifugation, the ethanol was aspirated and the pellet was air-dried for 5 minutes and heated at 65°C for 1-2 minutes to aid in the pellet dissolution. RNA pellets were then suspended in DNase-/RNase-free water. Total RNA concentration and purity was assessed using a Nanodrop 2000 (Thermo Fisher Scientific, Waltham, MA). RNA was stored at −80°C.

### cDNA synthesis

DNA was synthesized using the Thermo Fisher High Capacity Reverse Transcription kit according to manufacturer’s instructions (Thermo Fisher Scientific, Waltham, MA). Briefly, a mastermix containing reverse transcriptase (RT), RT buffer, dNTPs, random primers, and water was added to 30-50 μg of RNA. Samples were run on the Thermo Fisher QuantStudio 3 PCR machine using the following protocol: 25°C 10 minutes, 37°C 120 minutes, 85°C 5 minutes, 4°C hold.

### Quantitative RT-PCR

Quantitative PCR was performed using Thermo Fisher’s QuantStudio 3 and TaqMan technology (Thermo Fisher Scientific, Waltham, MA). TaqMan assays and the TaqMan Fast Advanced Mastermix were used according to manufacturer’s protocol. Briefly, mastermix, assays, water and cDNA were placed in individual wells of an optical 96 well reaction plate. Samples were run using the following parameters: 50°C 2 minutes, 95°C 10 minutes, 40 cycles of 95°C 15 seconds and 60°C 1 minute, and 4°C hold. TaqMan assays used are outlined in **Table S1**.

Differential gene expression was calculated using the comparative threshold cycle method (35). As described by Schmittgen and Livak, since all samples came from different animals or human patients, there is no means to justify which positive sample is compared with which negative sample and therefore the 2^-ΔΔCq^ method of relative gene expression quantification could not be used (35). Here, the mean ± standard error was calculated as individual data points using 2^-ΔCq^, where ΔC_q_ = C_q_ (gene of interest) – C_q_ (reference gene; *GAPDH*) (35). Where feasible, fold-changes in gene expression were calculated using these 2^-ΔCq^ values. Several C_q_ values, particularly in control non-tumor-bearing samples, were classified as undetermined and therefore relative gene expression could not be quantified for all groups (**Fig S1–S3**). Individual data points are presented in graphical form on a log_2_ scale, i.e. displaying the –ΔC_q_ values.

### *In vivo* model

All animal studies were approved by and performed in compliance with the Institutional Review Board for the Animal Care and Facilities Committee at Rutgers University and institutional guidelines on animal handling. Female athymic nude mice were purchased from Charles River Laboratories (Fairfield, NJ) and were received between 4-5 weeks of age and allowed to acclimate for one week before initiation of study. They were housed in sterile disposable cages and provided food and water *ad libitum*.

In order to assess longitudinal changes reflected in sEV profiles, 1×10^5^ SKOV-3 cells were injected intraperitoneally and allowed to grow for 3 weeks. SKOV-3 cells were previously tagged with red-fluorescent protein (RFP) allowing for weekly imaging validation of tumor growth prior to blood collection. Whole body fluorescence imaging was performed with the Bruker In Vivo MS FX PRO system (Carestream, Woodbridge, CT). Image analysis was performed analyzing pixel fluorescence intensity using ImageJ. Mice were separated into tumor-bearing and non-tumor-bearing groups (3-6 mice in each group), as well as time point groups: 5-7 days post injection of tumor cells, 10-15 days post injection, 20-25 days post injection. All groups underwent various imaging procedures for tumor identification. At each time point, animals from that group were euthanized according to university and protocol guidelines and whole blood was extracted by cardiac puncture. Blood was pooled between 2-3 animals in order to have a sufficient starting working volume for RNA extraction from the sEVs. Reported numbers refer to groups of pooled samples (n=3 means 3 separately pooled groups of 2-3 samples each, total 6-9 animals). Animals in the last time point group were euthanized at the compassionate endpoint after tumor burden correlated with ascites accumulation. Within 4 hours of collection, whole blood was separated by centrifugation at 3000 x g for 10 minutes into plasma, buffy layer, and red blood cells. Plasma was separated and stored at −80°C until further processing.

Ascites was collected post-mortem with a 25-26G needle injected into the peritoneal cavity. Some animals had thick mucousy ascites, requiring the opening of the peritoneal cavity to successfully collect the ascites. For early stage collection, 1 ml of sterile PBS was injected into the peritoneum and then extracted to obtain a “peritoneal wash” comparative to the ascites collected at later stages. Peritoneal wash from 2 animals was combined to obtain enough sample from which to extract sEVs, achieving n=3 from 6 animals. Sufficient volumes of ascites were collected from individual animals in the final week, allowing for a higher n number.

### Human serum collection and processing

Blood from ovarian cancer patients was collected in BD Vacutainer tubes (BD 367988) from Biorepository Services at the Rutgers Cancer Institute of New Jersey, under a Rutgers Institutional Review Board exemption. Following centrifugation (1000 x g, 10 minutes, room temperature), serum aliquots were immediately frozen at −80°C until use. Normal de-identified human whole blood and serum were obtained from Innovative Research, Inc. Whole blood was collected and processed by the company as outlined in their protocol. Briefly, blood was centrifuged at 5000 x g for 10 minutes. Supernatant was collected and using a plasma extractor into a separate transfer bag, where it was allowed to clot at room temperature up to 48 hours. Supernatant was then centrifuged at 5000 x g for 20 minutes at 4°C. Serum was separated and stored at 4°C until shipped.

### Bioinformatic analysis of gene signature correlation with patient outcomes

Assessment of the clinical relevance of the 10-gene panel was performed by analyzing gene expression in multiple existing databases. We used the OvMark algorithm (36, 37) which was designed to mine multiple international databases (14 datasets) of ovarian cancer patient outcomes for gene expression correlation (about 17,000 genes). Datasets that were used included GSE26712, GSE13876, GSE14764, GSE30161, GSE19161, GSE19829, GSE26193, GSE18520, GSE31245, GSE9899, GSE17260, GSE32062, TCGA, and an in-house dataset. Each of the 10-genes was evaluated individually in relation to progression free survival. Multiple parameters were utilized in the analysis, including a median expression cutoff, histology (serous and endometrioid), and disease stage. The cutoff level indicated that the median expression level of gene was used to determine high and low expression groups. Data was expressed in terms of a hazard ratio, which used Cox regression analysis to establish survival analysis. If the hazard ratio was greater than 1 then it corresponded to poor outcomes, with increasing numbers indicating the degree to which that poor outcome was expected in the high expression group. A hazard ratio less than one correlated with a good prognosis. A log-rank p value was determined for the differences between the high and low expressions of the gene.

### Statistics analysis

Statistical analysis was performed using GraphPad Prism 8 software. For qPCR data, statistical analysis was performed on ΔC_q_ values. Data were analyzed using one-way ANOVA followed by Tukey’s post-hoc test (comparison of 3 or more groups) or unpaired two-tailed t-test for comparison of two groups, with p <0.05 considered statistically significant.

## Results

### Genetic biomarkers correlate with poor clinical outcomes in ovarian cancer

In order to address the effectiveness of EV-derived gene signatures to predict disease progression we first evaluated the correlation of a known 10-gene panel (**Table 1**), selected from multiple datasets (12, 28–32), to disease prognosis. The correlation of the expression of the 10 genes in the panel to disease-free survival was assessed in multiple patient datasets (**Table S2**) using OvMark, a bioinformatics tool developed by Molecular Therapeutics for Cancer, Ireland (MTCI) (36, 37). Individual genes were added to the algorithm to determine patient outcomes related to overexpression of these genes. The algorithm determines a “hazard ratio” using a Cox regression analysis which associates increased gene expression with either a poor outcome (>1) or a good outcome (<1). Differences in high expression and low expression were assessed in relation to disease free survival using Kaplan-Meier estimates and log-rank p values. Correlation of disease-free survival was assessed for degree of overexpression, histological subtype, and disease stage. Eight genes (*THBS1, TIMP3, LOX, ACTB, COL5A1, AEBP1, COL11A1*, and *POSTN*) of the 10-gene panel showed that high expression had a significantly greater correlation with disease-free survival than low expression levels (**Fig 1, Table S3**). *TIMP3* and *POSTN* showed the greatest difference between high (black line) and low (gray line) expression levels. Neither *NECTIN4* nor *SNAI1* showed a significant difference between high and low expression and prognosis. Increased expression of all of the selected genes except *NECTIN4* and *SNAI1* showed significant correlation with poor clinical outcomes in patients with serous ovarian cancer, but not with the endometrioid subtype (**Table S4**). Two genes, *LOX* and *THBS1*, did show correlation with a positive clinical outcome in endometrioid ovarian cancer despite a lack of significance between the high and low expression levels (**Table S4**). Differences in gene expression were also assessed at varying tumor grades (**Table S5**). Grade 3 tumors showed the greatest difference between high and low expression levels, while both *POSTN* and *TIMP3* also showed significant differences in grade 1 (**Table S5**). *NECTIN4* and *SNAI1* both expressed hazard ratios that were less than 1 in grade 3, while *NECTIN4* had similar results in grade 2 tumors (**Table S5**), signifying that while there was not a significant correlation the trend suggests a need for further research into their potential as positive prognosticators.

**Figure 1.**
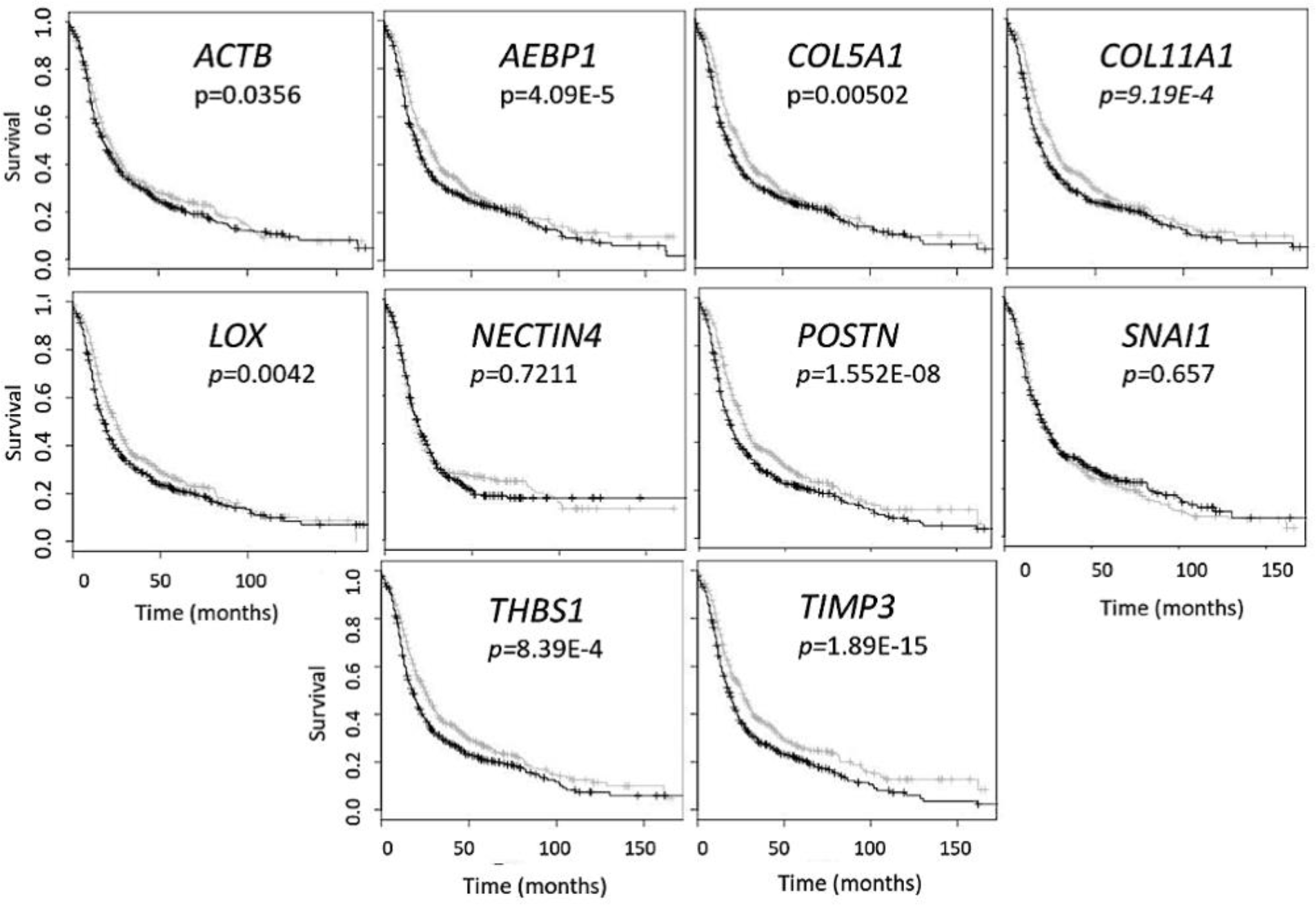
Gene panel expression correlates with disease-free survival in ovarian cancer patients. Kaplan-Meier curves show the difference between high expression (black line) of the individual gene and low expression (grey line). The OvMark algorithm used Cox regression analysis and log-rank p values to determine the difference in expression levels. *NECTIN4* and *SNAI1* were the only two genes to not show a significant difference between high and low expression of the disease with regards to disease-free survival.

**Table 1.** Genetic biomarkers of ovarian metastasis

| Gene name | Overall function | Ovarian cancer function | References |
| --- | --- | --- | --- |
| *ACTB* | Cell motility, stabilization, intercellular signaling | Invasion, metastasis | (38) |
| *AEBP1* | Transcriptional repression, cell differentiation | Regulate proliferation, increased NF-κB signaling, collagen remodeling | (39) |
| *COL5A1* | Extracellular matrix (ECM) structure, binds DNA | Proliferation, migration, chemoresistance | (40) |
| *COL11A1* | ECM structure and binding | Cell invasion, tumor formation | (41) |
| *LOX* | Collagen crosslinking, ECM remodeling | Tumor cell invasion, collagen crosslinking | (42) |
| *NECTIN4* | Cell adhesion | Epithelial-mesenchymal transition (EMT), adhesion, migration, proliferation | (43) |
| *POSTN* | Adhesion and migration | Cancer stem cell maintenance and metastasis, EMT | (44) |
| *SNAI1* | EMT, cell migration, transcription repression | EMT, invasion, proliferation | (45) |
| *THBS1* | Binds extracellular proteins, regulates intercellular interactions | Regulate growth and adhesion, migration of tumor cells | (46) |
| *TIMP3* | Inhibits matrix metalloproteinases | Tumor suppressor, inhibits angiogenesis | (47) |

### Plasma-, serum-, and ascites-derived extracellular vesicles conform to size characteristics of small extracellular vesicles (sEVs)

While standardized protocols and guidelines for the isolation and purification of extracellular vesicles (EVs) are still under debate, particle sizing is a commonly accepted method of vesicle classification (48, 49). Based on guidance issued by the International Society for Extracellular Vesicles, current optical measurements such as dynamic light scattering and nanoparticle tracking analysis (NTA) are used for characterization and quantification of extracellular vesicles (49). We characterized small extracellular vesicles (sEVs) isolated from plasma/serum and ascites using a commercial polymer precipitation method (ExoQuick). Isolated sEVs had a mean size of 126.6 nm (mean range of 78.5-157.5 nm) and a mode size of 96.5 nm (mode range of 63.1-119.4 nm) and representative histograms of sEV sizes from mouse plasma, mouse ascites, and human serum are as shown in **Figure 2a-c** respectively. **Figure 2d** shows a representative snapshot of sEVs measured using Nanosight NS300. The sizes were in accordance with acceptable ranges for sEVs (24, 49). Specifically, mouse plasma had a mean size of 125±3 nm and mode size of 96.5±3 nm (**Fig 2e**). Mouse ascites-derived vesicles were slightly larger with a mean size of 145.1±6.7 nm and a mode of 114.3±1.3 nm (**Fig 2e**). Vesicles isolated from human serum-derived vesicles had a mean size of 121.5±4.9 nm and mode size of 97.1±4.5 nm (**Fig 2e**). Vesicle size was similar independent of source of material (p = 0.106, one-way ANOVA for mean size, p = 0.175, one-way ANOVA for mode size), indicative of consistency in extracellular vesicle isolation results and comparable EV populations between biofluid sources and tumor conditions evaluated in this study.

**Figure 2.**
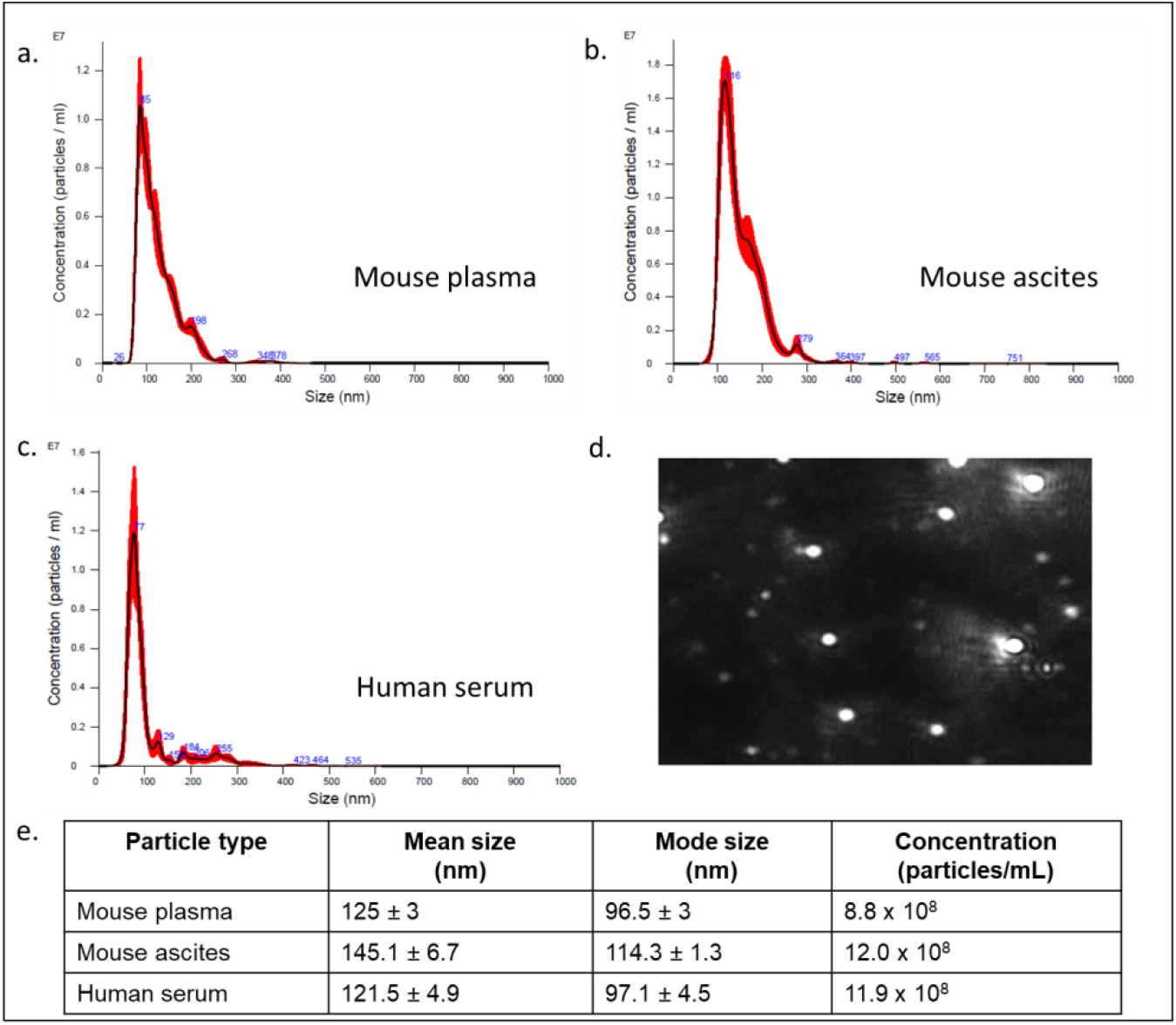
Small extracellular vesicle (sEV) isolation from mouse plasma, ascites, and human plasma shows similar sized populations independent of biofluid. Representative sizing of vesicles isolated from **a**. mouse plasma, **b**. mouse ascites, and **c**. human serum. **d**. Representative image of mouse plasma vesicles on the Nanosight N300. **e**. sEV from plasma from mice had an average mode size of 96.5±3 nm and concentration of 8.8×10^8^ particles/ml. sEV from ascites from mice had an average mode size of 145.1±6.7nm and a concentration of 12.0×10^8^ particles/ml. sEV from human serum had an average mode size of 121.5±4.9nm and a concentration of 11.9×10^8^ particles/ml.

### Genetic expression levels change with longitudinal tumor development and progression in a mouse model of ovarian cancer

The challenge in ovarian cancer is early detection of disease, as the current screening tools are either not sensitive enough or invasive screening techniques are usually not employed at the earlier asymptomatic stages of the disease. Existing screening tools that are invasive and require biopsied specimens do not allow for longitudinal monitoring of a probable genetic signature that can predict metastatic potential and tumor progression. The advantage of the sEV-based liquid biopsy tool would be the ability of sEVs to package the genetic information from the tumor and its microenvironment allowing for its preservation once it is in the peripheral circulation. In order to assess the potential of sEVs as screening tools we determined the validity of the 10-gene signature to correlate with tumor progression in a SKOV-3 (human epithelial ovarian cancer cell line)-derived mouse model of ovarian cancer metastases. SKOV-3 cells were injected into the peritoneal cavity of athymic nude mice, and plasma was collected from animals euthanized at weekly intervals. Tumor growth in the peritoneal cavity was validated by fluorescent imaging (**Fig 3a-c**). sEVs were extracted from the collected plasma and the 10-gene signature was evaluated. Of the 10-gene signature, seven genes were expressed at quantifiable levels as determined by qRT-PCR.

**Figure 3:**
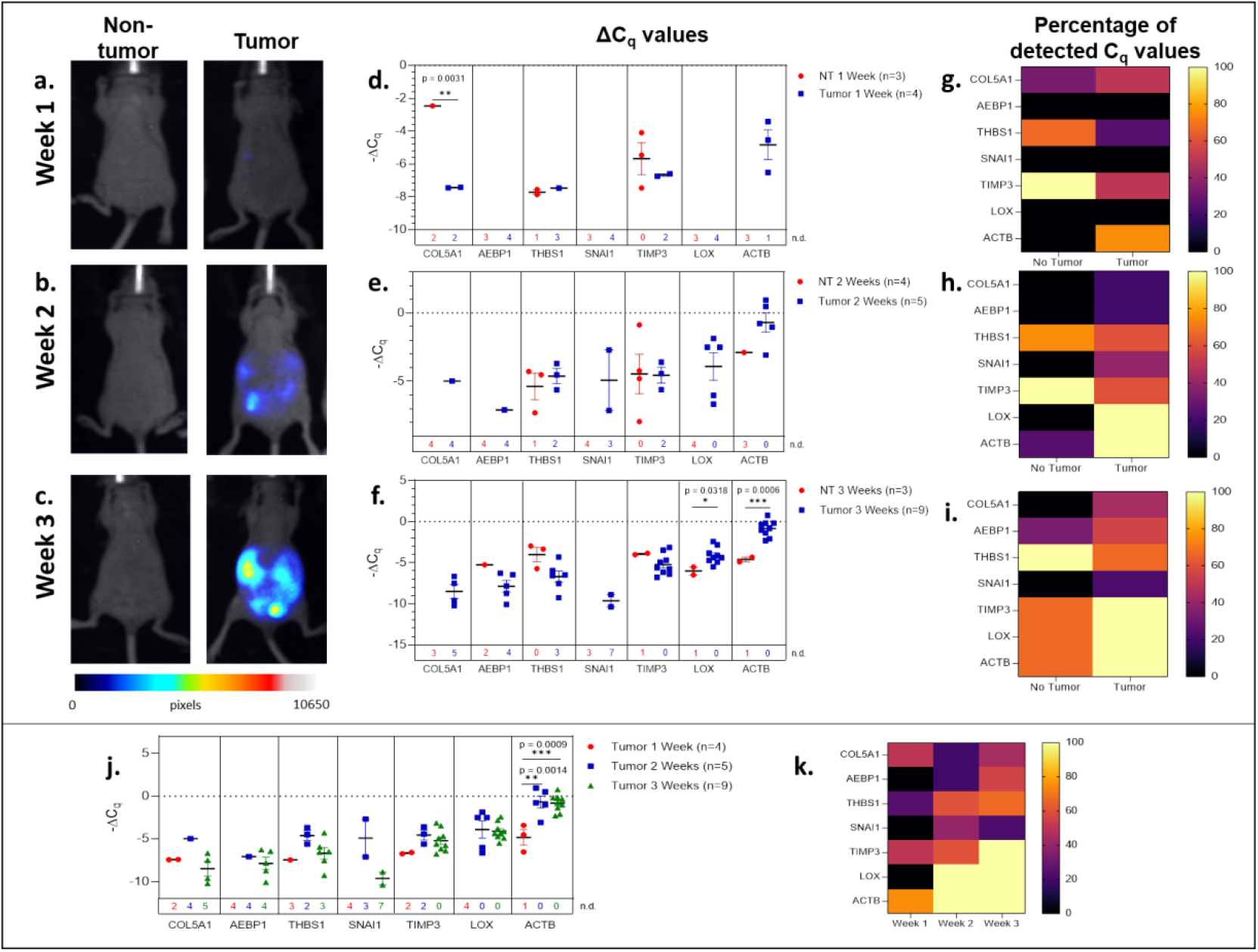
Plasma-derived sEV gene expression in a mouse model of ovarian cancer. Representative fluorescent imaging of SKOV-3/RFP cells in tumor-bearing and non-tumor-bearing mice in **a.** Week 1, **b**. Week 2, and **c.** Week 3. Scatter plots of ΔC_q_ values at **d.** Week 1 for tumor- (blue, n=4) and non-tumor-bearing samples (red, n=3), **e**. Week 2 for tumor- (blue, n=5) and non-tumor-bearing samples (red, n=4), and **f.** Week 3 for tumor (blue, n=9) and non-tumor-bearing samples (red, n=3). Heat maps showing the percentage of detected C_q_ values at **g.** Week 1, **h.** Week 2, and **i.** Week 3. **j**. Scatter plot of ΔC_q_ values for tumor-bearing samples over Weeks 1-3 of tumor development; **k**. Heat map showing the percentage of detected C_q_ values in tumor-bearing samples for Weeks 1, 2, and 3. p values for unpaired two-tailed t-test are labeled in the graphs. The number of non-detected (n.d.) C_q_ values in each experimental group are listed underneath the corresponding scatter plots. Heat maps in g, h, i, and k indicate the absence/presence of the target gene (percentage of detected C_q_ values) in each experimental group.

We initially compared plasma-derived sEV gene expression in tumor-bearing animals to non-tumor-bearing controls during weeks 1, 2, and 3 of tumor development. As shown in **Figures 3d and 3e**, there was minimal or no expression of the 10-gene signature in non-tumor-bearing mice during weeks one and two of tumor development, which precluded quantification of differential gene expression. However, this absence of target gene expression in control mice (highlighted by the number of non-detected C_q_ values underneath the scatter plots and **Fig S1**) compared to the presence of the target gene in tumor-bearing animals does indicate an increase in gene expression. During week 3 of tumor development, non-quantifiable increases in gene expression of tumor-bearing sEVs were observed in *COL5A1* and *SNAI1* (absence vs. presence of target gene C_q_ values), and significant quantifiable increases in gene expression were observed in *LOX* (*p* = 0.0318, 4.37-fold change) and *ACTB* (*p* = 0.0006, 16.3-fold change) (**Fig 3f**). The corresponding heatmaps showing the percentage of detected C_q_ values for weeks 1-3 are as shown in **Figure 3g-i** respectively.

There is a correlation between gene signature and tumor progression suggesting the potential for sEV-based liquid biopsy as a longitudinal monitoring tool in ovarian cancer. As shown in **Figure 3j**, *ACTB* expression significantly increased after the first week of tumor progression and remained elevated until the end of the experiment (*p* = 0.0007, One-way ANOVA; Tukey’s post-test: Week 1 vs Week 2: *p* = 0.0014, Week 1 vs Week 3: *p* = 0.0009). This corresponds to an 18.4-fold increase in *ACTB* expression between weeks 1 and 2 of tumor development. The absence/presence of target gene expression (i.e. undetermined vs. measured C_q_ values) is another indicator of increasing gene expression. Expression of genes such as *AEBP1, SNAI1*, and *LOX* similarly increased after the first week of tumor progression, although these changes could not be quantitatively compared to week 1 due to undetermined C_q_ values at this time point (**Fig 3d-f**).

### Select genes were differentially expressed in sEVs isolated from human serum based on tumor presence, histopathological characterization, and TNM staging

In order to determine the fidelity of the signature from mouse ovarian cancer studies, we evaluated the expression of the 10-gene panel in a small cohort of patient samples. We determined the ability of the 10-gene signature to distinguish between tumor and non-tumor/normal human samples, and within the tumor samples stratified by metastasis, histological subtypes, or TNM (tumor, node, and metastasis) staging. To this end, we compared serum-derived sEVs from eleven patients with ovarian cancer and three cancer-free patients. Collected samples came from different histological subtypes and varying TNM stages (**Table S6**) and the 10-gene signature was evaluated using quantitative RT-PCR. Three genes within the 10-gene signature (*ACTB, THBS1*, and *TIMP3*) were found to be expressed at quantifiable levels in the collected patient samples. The most notable difference was seen when comparing metastatic vs non-metastatic tumors, where the samples from patients with metastatic tumors had significantly higher expression of *THBS1* compared to those from patients with non-metastatic tumors (*p* = 0.0192, 4.53-fold change) (**Fig 4a**). However, when these patient samples were scored based on TNM staging, due to small sample sizes and non-quantifiable levels of mRNA (i.e. undetermined C_q_ values) no significant differences were found (**Fig 4b**). Although trends indicate an increase in *ACTB* and *THBS1* expression in ovarian cancer patients compared to healthy control subjects, differences in ΔC_q_ values were not significantly different due to low sample numbers (**Fig 4c**). The limited expression of these genes in the healthy control subjects hindered robust statistical analysis of differential gene expression. As mentioned earlier, the absence/presence of target gene expression (i.e. undetermined vs. measured C_q_ values; **Fig S2**) is another indicator of increasing gene expression in the tumor-bearing serum samples (50, 51). As shown in **Table S6**, the patient samples were also stratified into 4 different histological subtypes: serous, clear cell, endometrioid, and mixed. No significant differences were found in sEV gene expression between these histological subtypes, likely due to small sample sizes and non-quantifiable levels of mRNA (i.e. undetermined C_q_ values) for several of these samples (**Fig 4d**). Due to low sample sizes and limited expression of the selected genes in the healthy control samples, it was difficult to evaluate the ability of serum-derived sEVs to indicate ovarian tumor presence in human samples in this study.

**Figure 4.**
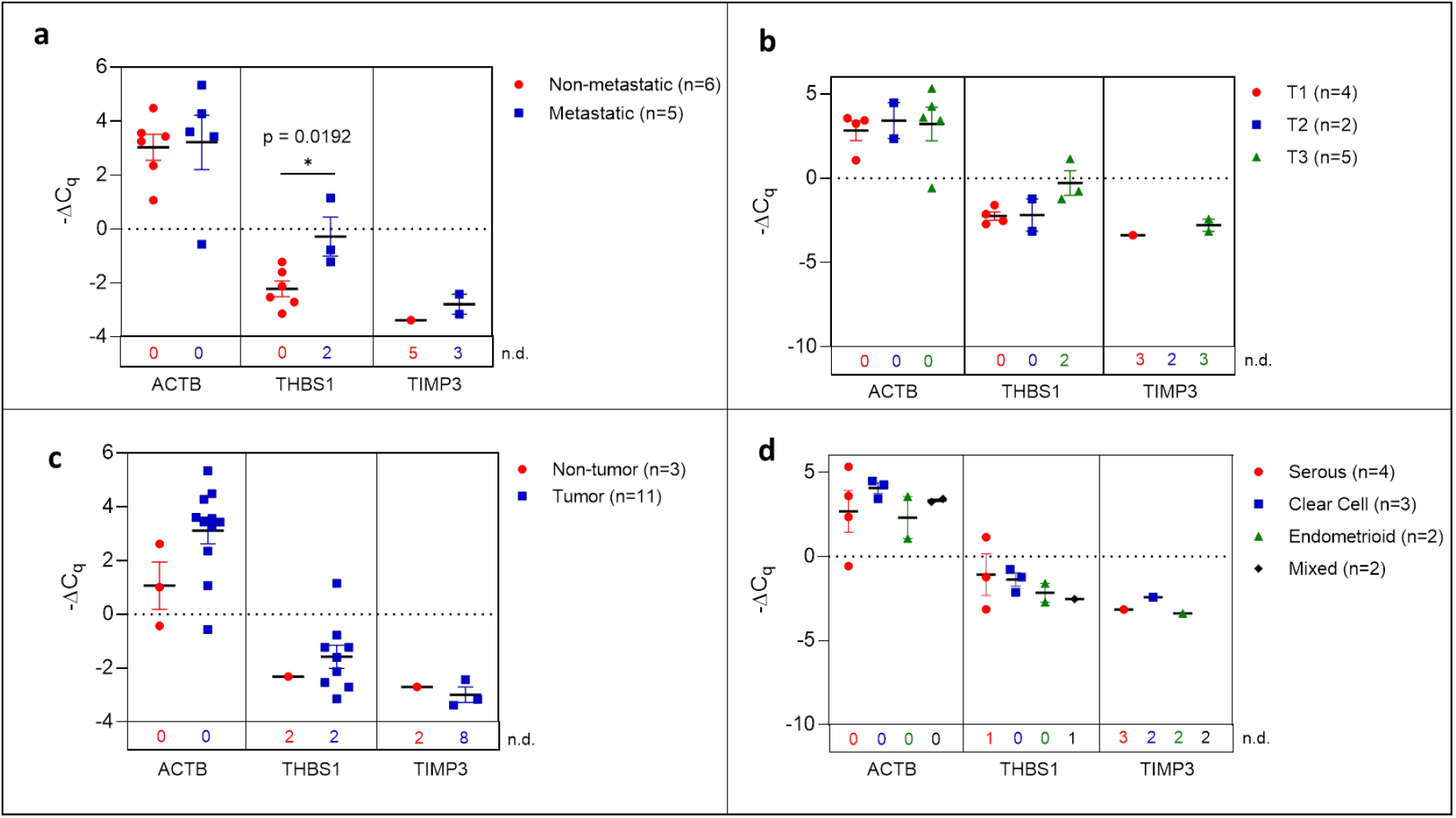
Plasma-derived sEV gene expression in human samples. Scatter plots of ΔC_q_ values comparing **a.** serum from metastatic and non-metastatic samples, **b**. samples stratified by TNM staging **c**. tumor-bearing and non-tumor-bearing samples, and **d**. samples stratified by histology. P values indicated were determined via unpaired two-tailed t-test. The number of non-detected (n.d.) C_q_ values in each experimental group are listed underneath the corresponding scatter plots.

### Metastasis markers identified in sEVs extracted from ascites

A hallmark of advanced ovarian cancer is the presence of ascitic fluid in the peritoneal cavity and palliative therapy often requires repetitive drainage of this fluid (52) making this a possible analyte for liquid biopsy. In the SKOV-3 peritoneal metastasis mouse model, we determined the 10-gene signature trend in correlation to disease progression from ascites-derived sEVs. During the first week of tumor development and in non-tumor-bearing mice, ascites was not present. In order to assess sEV profiles at early stages, PBS was injected into the peritoneum and extracted as a peritoneal wash containing sEVs. This was then compared to the ascites collected during later stages of disease progression.

Initially, ascites-derived sEV gene expression in tumor-bearing animals was compared to non-tumor-bearing controls during longitudinal tumor progression. Due to the absence of measurable ascitic fluid and similar to plasma-derived sEVs, there was limited or no expression in non-tumor-bearing mice during weeks one (**Fig 5a**) and two (**Fig 5b**). The absence of the target gene (undetermined C_q_ values) in control animals compared to the presence of the target gene in tumor-bearing animals again indicates a non-quantifiable increase in gene expression (**Fig S3**).**Figure 5b** illustrates this non-quantifiable increase in the expression of *COL5A1, AEBP1, SNAI1, LOX*, and *ACTB* in tumor-bearing mice after 2 weeks of tumor development. During week 3 of tumor development, non-quantifiable increases in gene expression were observed in *COL5A1, THBS1, COL11A1*, and *SNAI1*, and significant quantifiable increases in gene expression were observed in *AEBP1* (*p* = 0.0379, 46.5-fold change), *LOX* (*p* = 0.0017, 5.81-fold change), and *ACTB* (*p* < 0.0001, 22.6-fold change) (**Fig 5c**). This trend of increasing gene expression over the three weeks of tumor progression is also evident when comparing ΔC_q_ values of the tumor-bearing animals over time (**Fig 5d**). There were significant increases in *COL5A1* (*p* = 0.0332, one-way ANOVA), *TIMP3* (*p* = 0.0206, one-way ANOVA; Tukey’s post-test: *p* = 0.0248, Week 1 vs Week 3), *LOX* (*p* = 0.0037, one-way ANOVA; Tukey’s post-test: *p* = 0.0029, Week 1 vs Week 3), and *ACTB* expression (*p* = 0.0047, one-way ANOVA; Tukey’s post-test: *p* = 0.0035, Week 1 vs Week 3) at week 3 compared to week 1. These differences in ΔC_q_ values correspond to a 19.3-fold change in *COL5A1* expression, 61.3-fold change for *TIMP3*, 3.92-fold change for *LOX*, and 6.12-fold change for *ACTB*. The smaller fold-change increases in longitudinal expression of *LOX* and *ACTB* suggest that these two genes are expressed more strongly in ascites-derived sEVs at earlier time points compared to other genes in the 10-gene signature. The gene expression patterns seen in the ascites-derived sEV samples validate those seen in the plasma-derived sEV samples, and further demonstrate the potential for sEVs to distinguish between tumor-bearing and non-tumor-bearing samples.

**Figure 5.**
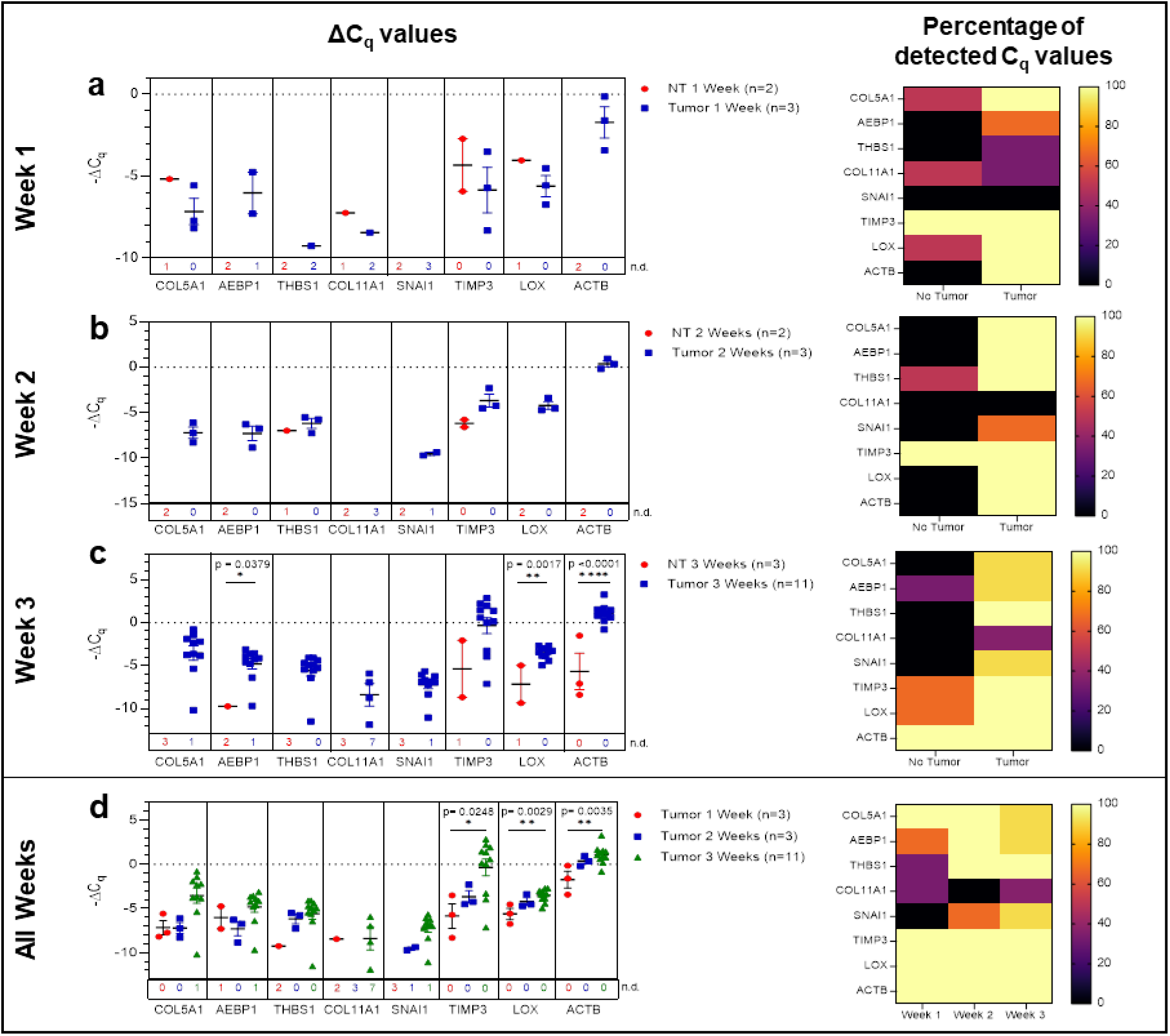
Ascites-derived sEV gene expression in a mouse model of ovarian cancer. Scatter plots of ΔC_q_ values and heat maps showing the percentage of detected C_q_ values at **a.** Week 1, tumor- (blue, n=3) and non-tumor-bearing samples (red, n=2); **b.** Week 2, tumor- (blue, n=3) and non-tumor-bearing samples (red, n=2); **c.** Week 3, tumor- (blue, n=11) and non-tumor-bearing samples (red, n=3); **d.** over Week 1 (red, n=3), Week 2 (blue, n=3), and Week 3 (green, n=11) of tumor development. p values for unpaired two-tailed t-test are labeled in the graphs. The number of non-detected (n.d.) C_q_ values in each experimental group are listed underneath the corresponding scatter plots. Heat maps indicate the absence/presence of the target gene (percentage of detected C_q_ values) in each experimental group.

### Correlation of genetic signature between tumor-derived sEVs from the niche and peripheral circulation

One of the major barriers in the success of liquid biopsy tools as diagnostics is the lack of standardization and reproducibility (49, 53, 54). In order to establish a liquid biopsy tool that can prognosticate disease progression, it is important to determine the correlation between gene signature at tumor site and in peripheral circulation. Hence, we determined if the TME genetic signature would correlate with the signature obtained from peripheral circulation. Towards this, we compared gene expression of the ascites-derived sEVs from the peritoneal cavity to that of the plasma-derived sEVs (**Fig 6**), demonstrating a concordance in expression patterns overall.

**Figure 6.**
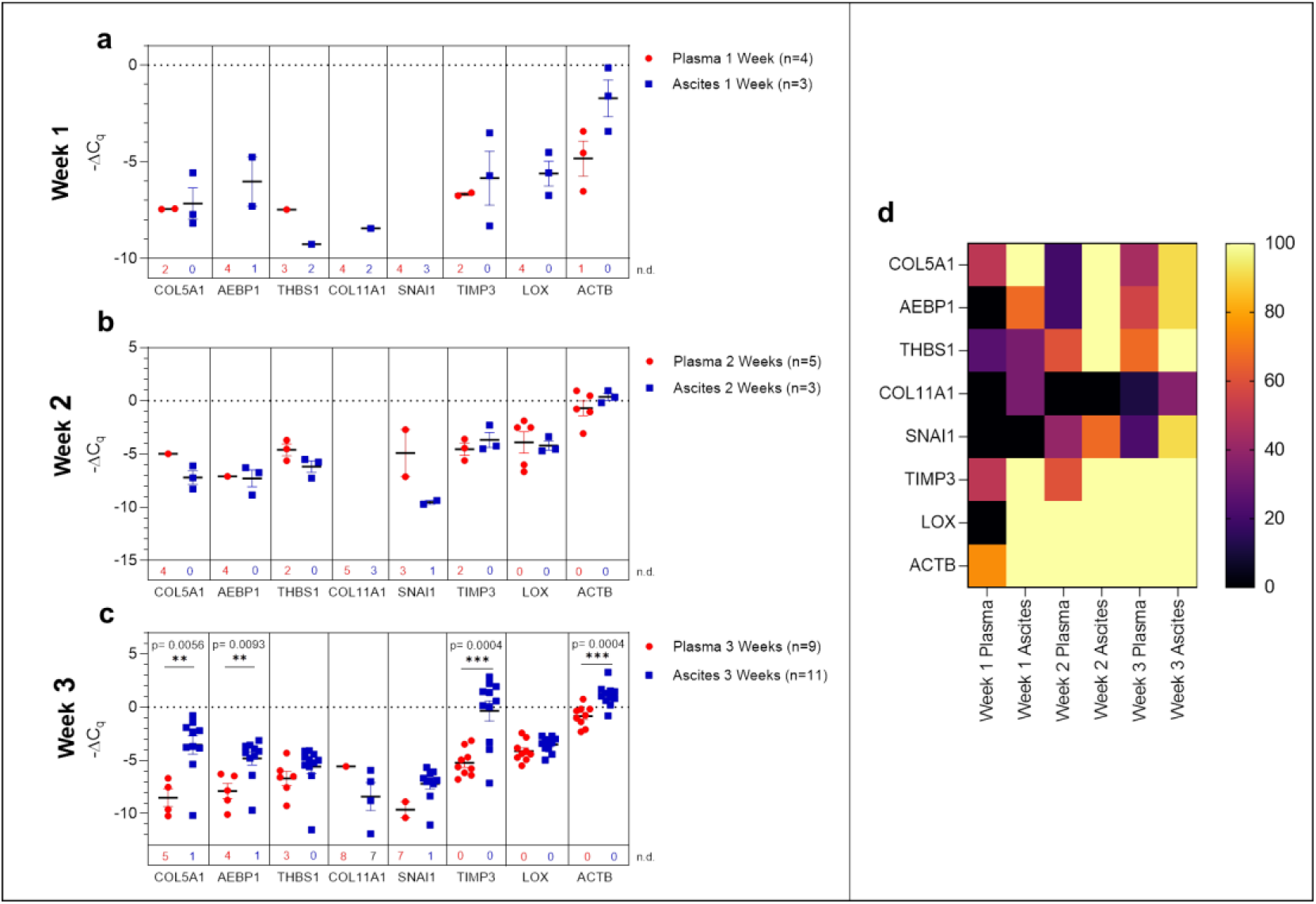
Comparison of plasma-derived and ascites-derived sEV gene expression in a mouse model of ovarian cancer. Scatter plot of ΔC_q_ values for plasma-derived and ascites-derived sEVs at: **a.** Week 1 plasma (red, n=4) and ascites (blue, n=3); **b.** Week 2 plasma (red, n=5) and ascites (blue, n=3); **c.** Week 3 plasma (red, n=9) and ascites (blue, n=11). **d.** Heat map showing the percentage of detected C_q_ values over Weeks 1-3 indicates the absence/presence of the target gene in each experimental group. p values for unpaired two-tailed t-test are labeled in the graphs. The number of non-detected (n.d.) C_q_ values in each experimental group are listed underneath the corresponding scatter plots.

During early tumor progression (weeks 1 and 2 of development), there was no significant difference in plasma-derived and ascites-derived sEV gene expression, however, differences in expression of certain genes (*AEBP1, COL11A1*, and *LOX*) could not be quantified at week 1 due to undetermined C_q_ values in the plasma samples (**Fig 6a and 6b**). In the final week of tumor progression in the mice with the expected late-stage development of ascites, gene expression was overall higher in the ascites-derived samples, with significant increases in *COL5A1* (*p* = 0.0056, 43.1-fold change), *AEBP1* (*p* = 0.0093, 8.54-fold change), *TIMP3* (*p* = 0.0004, 58.4-fold change), and *ACTB* (*p* = 0.0004, 3.98-fold change) (**Fig 6c**). The data suggest that sEVs sampled from ascitic fluid are likely a stronger indicator of tumor presence compared to sEVs sampled from plasma, though both show upregulation of genes in the 10-gene signature panel (**Fig 6d**) that correspond to longitudinal changes in tumor progression.

## Discussion

Ovarian cancer treatment has been a challenge due to its asymptomatic nature in early stages of the disease leading to eventual detection at advanced stages, which then results in low survival rates (1, 2, 14–16). Current screening and monitoring tools for ovarian cancer lack specificity and often lead to false positives, which require invasive follow-up biopsies (4–7, 13). There is a need for non-invasive monitoring tools that can perform with more sensitivity than the current tools, transvaginal ultrasound and CA-125 testing.

The emerging era of genomics is transforming the field of oncology by allowing for development of new diagnostics and therapeutics that tailor to specific tumor types and stages (55). Liquid biopsy is the next generation diagnostic that integrates genetic signatures for disease profiling. To date, several liquid biopsy-based companion diagnostics have been approved by the FDA (56). While most of the approved diagnostics are typically based on circulating cell-free DNA, the use of EVs and exosomes as messengers of tumor presence (57) in order to improve diagnostic abilities is actively being explored. In 2019, the first diagnostic tool to employ EVs in clinical diagnostics, Bio-Techne’s ExoDx Prostate IntelliScore (EPI) test, was given FDA Breakthrough Device Designation, and is currently in use in the clinic (26). Circulating sEVs have the potential to address the weaknesses of tissue biopsies to monitor tumor progression and changes longitudinally. This can be applied in diagnostics (58), prognostics (59), and therapeutics (60), making it a crucial technology to advance (25, 61).

Our goal with this research was to establish a tumor-derived EV-based genetic signature that originated from gene expression analysis patient datasets of ovarian cancer. Peritoneal spread of metastatic lesions requires tumor cells to escape from the primary tumor, disseminate through the peritoneal cavity, adhere and invade into the peritoneal lining and then establish lesion growth (62). Each step is characterized by different molecular changes. The extracellular matrix (ECM) is a key component of the TME, and undergoes remodeling during many of the stages of metastasis establishment (63–65). This remodeling plays a key role particularly in the development and progression of many epithelial cancers including ovarian cancer (33, 34, 66–68). Our 10-gene signature, which overlaps with the signature elucidated by Cheon et al., was thus focused on collagen remodeling genes that are implicated in invasion and metastases in cancer (34, 67, 68). Through bioinformatic analysis using published datasets such as that of Cheon et al. (28), a ten gene signature that was overexpressed in ovarian cancer (28, 69) and was focused on collagen remodeling (**Table 1**) was selected. An established OvMark (36, 37) derived analysis of the 10-gene signature showed a clear association of eight of the ten genes to expression-based disease prognosis (**Fig 1**).

In this study, we explored the feasibility of a longitudinal sEV-based gene signature that would be predictive of metastasis progression in a mouse model of ovarian cancer. In this study, we isolated sEVs from both plasma and ascites from the mouse model and from human ovarian cancer patients’ serum. The sEVs were characterized based on NTA (**Fig 2**) and validated to conform to the acceptable size range for sEVs (49). We were able to show a quantifiable change in seven genes (*COL5A1, AEBP1, THBS1, SNAI1, TIMP3, LOX* and *ACTB*) of the 10-gene signature in plasma-derived sEVs longitudinally over the three-week period (**Fig 3**). Of the seven genes, *AEBP1* overexpression plays an important role in stimulating the crosstalk between the ECM and the pro-inflammatory NF-κB pathway inducing metastatic processes (29, 39), and *TIMP3*, a key regulator of ECM degradation that has been linked to a metastatic signature identifying aggressive tumors (12). *COL5A1, SNAI1, LOX*, and *ACTB* are mediators of ECM integrity (70). The most significant differences were observed in expression of *LOX*, a gene activated by hypoxia to enable invasive potential by crosslinking collagen (42) and *ACTB*, a gene very active in the metastatic process of epithelial to mesenchymal transition (EMT) and cell migration (38, 71, 72). We were able to observe significant quantifiable differences in only two of the seven genes mainly due to limited samples and a small animal cohort. Despite the lack of quantitative values to evaluate statistical significance, consistent undetermined values at initial time points (**Fig S1–S3**) represent a noteworthy change and the presence of a dynamic tumor environment being reflected in the sEVs. Our findings also extended to correlate the plasma-derived sEV signature obtained from our animal model to human serum-derived sEV from ovarian cancer patients. We found notable differences in *ACTB* and *THBS1* when compared to control serum, and in *THBS1* when comparing metastatic to non-metastatic patient samples (**Fig 4**). *THBS1* is known to play a role in cell-cell and cell-matrix interactions that are key for metastases progression to the peritoneal space in ovarian cancer (46). Given the limited nature of our patient cohort, future studies will be necessary to advance these findings to a larger cohort of samples. Additionally, sEV heterogeneity and contamination from non-tumor-derived sEVs also play a role in the fidelity and significance of the gene signature (73, 74). Future studies will focus on enrichment for cancer-specific sEVs to increase the sensitivity of the biomarkers for early detection, progression and metastasis.

Given the heterogeneous nature of the plasma-derived sEVs, in an effort to probe the reliability of the plasma-derived signature, we explored the significance of the 10-gene signature in disease progression from the TME. We isolated sEVs from ascites sampled from a mouse model of peritoneal ovarian cancer metastases and found a quantifiable change in seven genes (*COL5A1, AEBP1, THBS1, SNAI1, COL11A, LOX* and *ACTB*) of the 10-gene signature longitudinally over the three-week period (**Fig 5**). In addition to *LOX* and *ACTB*, which were highly significant in the plasma-derived sEV signature, there were also significant differences in *AEBP1* expression, with its pro-inflammatory stimulation of metastasis (29, 39). *COL11A1*, which was absent in plasma-derived sEVs but present in the ascites-derived signature, has been correlated with advanced disease stages (41).

When signatures from plasma- and ascites-derived sEVs were compared we found significant differences in expression of *COL5A1, AEBP1, TIMP3*, and *ACTB* at week 3 of tumor progression (**Fig 6**), suggesting that ascites-derived sEVs are likely a stronger indicator of tumor presence compared to plasma-derived sEVs. However, there were no significant differences in ascites vs. plasma-derived sEV expression of *THBS1, COL11A1, SNAI1*, and *LOX*. Further, plasma-derived sEV expression of *COL5A1* and *ACTB* could be used to indicate tumor presence when comparing tumor-bearing and healthy control samples at this time point. This pattern suggests that sEV contents in the periphery reflect changing molecular and functional states within the TME, enabling the use of plasma-derived sEVs as potential analytes for liquid biopsy to discern tumor progression.

While our focus was on demonstrating the plausibility of a sEV-based screening tool in a mouse model of ovarian cancer, several limitations of this study should also be acknowledged. One of the major limitations was while plasma would be the most suitable analyte for non-invasive screening, plasma-derived exosomal burden is low requiring larger plasma volumes. In our mouse model of ovarian cancer, this would have required a large cohort of animals per group for the longitudinal study. We had plasma volumes of <1ml even with pooled samples (n=2-3 animals) and this had an impact on exosomal burden and also the extracted RNA from this population. Correlation of our findings from the preclinical mouse model with plasma-derived sEVs from patients while encouraging, is still preliminary. Future studies will require larger patient cohort samples to further establish clinical validity of a sEV-based screening tool for ovarian cancer.

This study shows promise for the use of sEVs in cancer diagnosis and longitudinal disease monitoring in ovarian cancer. Liquid biopsies that use analytes such as sEVs carry tremendous clinical potential (75) but there is a need to validate our findings in larger patient cohorts towards developing a transformative non-invasive diagnostic with greater accuracy for early detection of ovarian cancer.

## Acknowledgements

The authors would like to thank the Rutgers University Molecular Imaging Core and Derek Adler for access to and assistance with the fluorescent imaging validation for this study. We also are appreciative to Arash Hatefi and Suzie Chen in the School of Pharmacy at Rutgers University for access to their Nanosight NS300 machine used in our NTA experiments.

## Author contributions

AG, PVM, and VG contributed to the conceptualization and design of the study. Investigation was performed by AG, NZ, JVS, JNS, BI, SG and VG. Formal analysis and project administration was performed by AG, NLF, and VG. PVM and VG were responsible for supervision. PVM, MK, and SKL contributed to resources and writing-review and editing. AG, NLF, and VG were responsible for visualization and writing-original draft preparation. All authors contributed to manuscript revision, read, and approved the submitted version.

## Data availability statement

The datasets generated for this study are available on request to the corresponding authors.

## Supplemental figure and table captions

**Supplemental figure 1:**
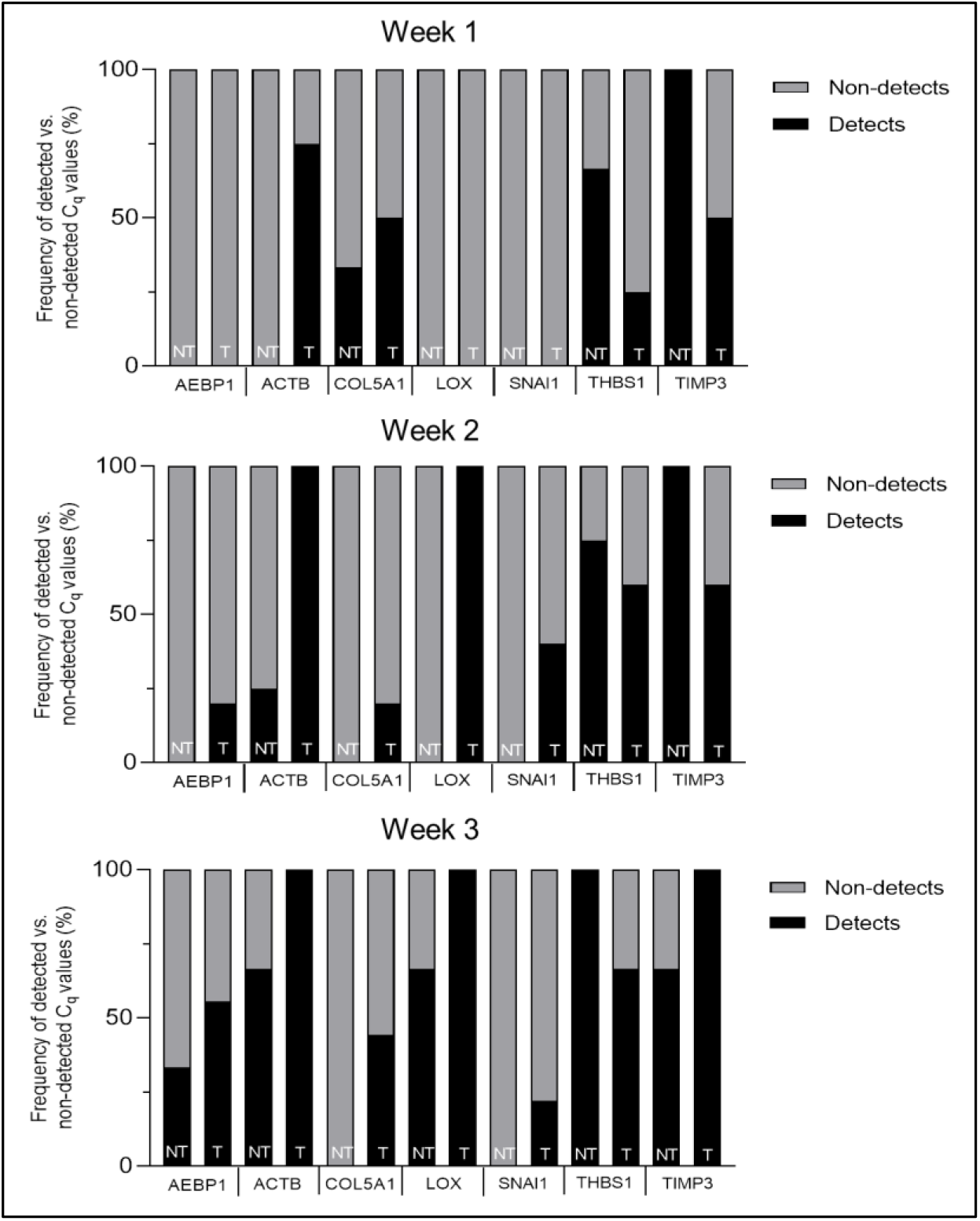
Frequency plots of detected vs. non-detected C_q_ values in plasma-derived sEV samples.

**Supplemental figure 2:**
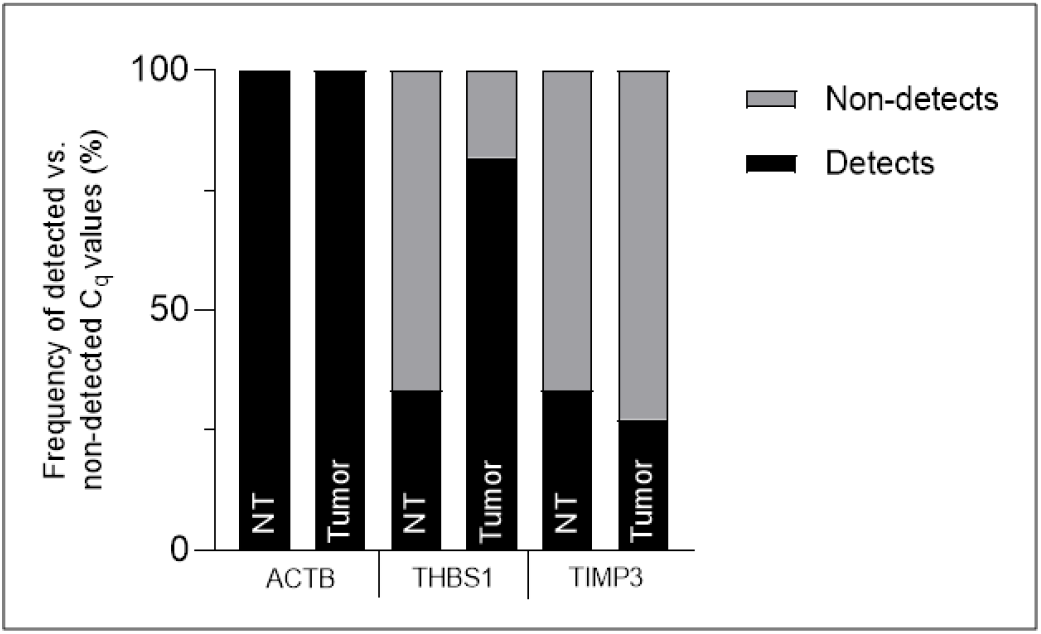
Frequency plots of detected vs. non-detected C_q_ values in human plasma-derived sEV samples.

**Supplemental figure 3:**
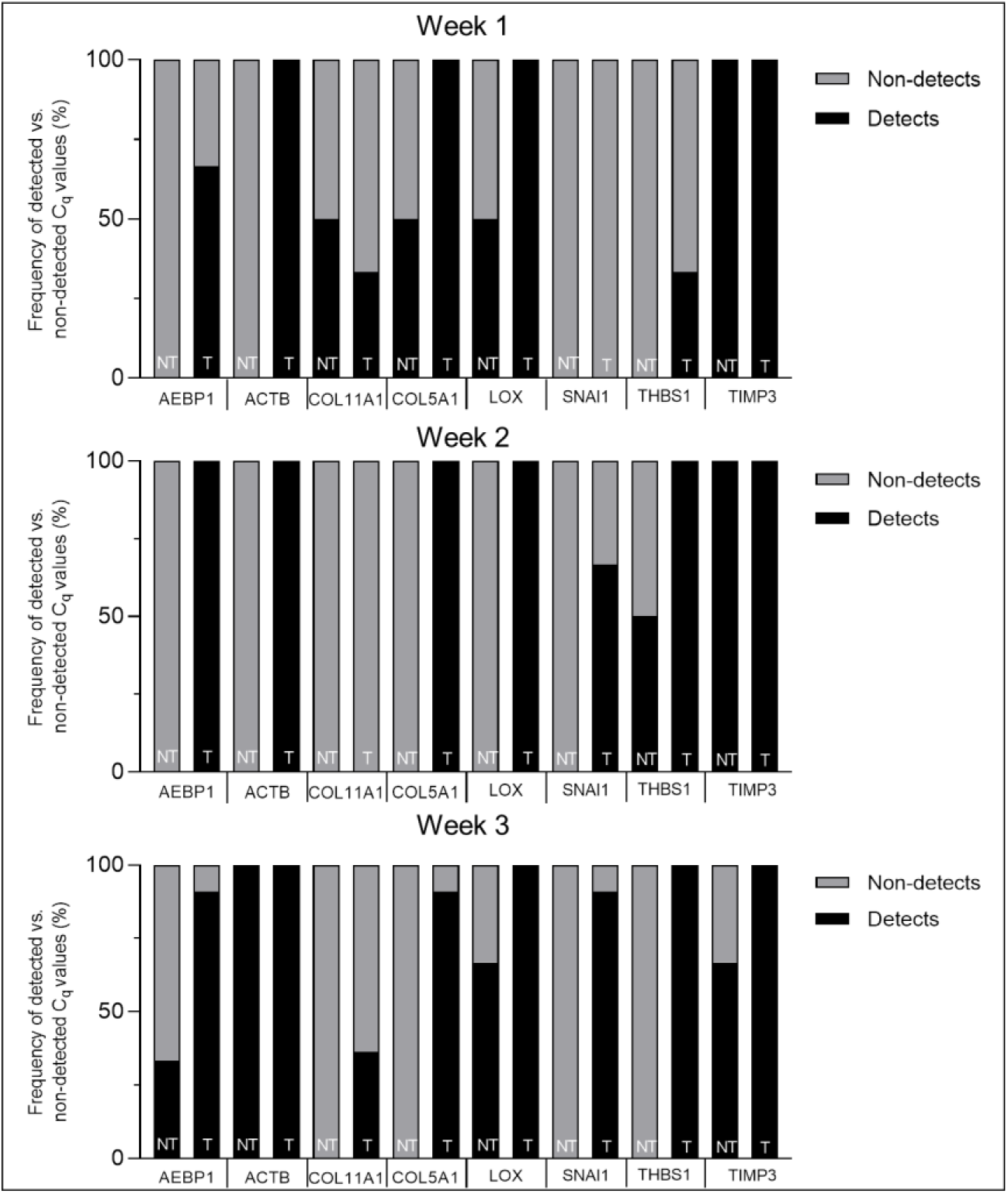
Frequency plots of detected vs. non-detected C_q_ values in ascites-derived sEV samples.

**Supplemental table 1:** TaqMan gene expression assays used for qRT-PCR purchased from Thermo Fisher.

| Gene Name | Gene expression assay ID |
| --- | --- |
| *ACTB* | Hs99999903_m1 |
| *AEBP1* | Hs00937468_m1 |
| *COL5A1* | Hs00609088_m1 |
| *COL11A1* | Hs01097664_m1 |
| *GAPDH* | Hs02786624_g1 |
| *LOX* | Hs00942480_m1 |
| *NECTIN4* | Hs00363974_m1 |
| *POSTN* | Hs01566750_m1 |
| *SNAI1* | Hs00195591_m1 |
| *THBS1* | Hs00962908_m1 |
| *TIMP3* | Hs00165949_m1 |

**Supplemental table 2:** Datasets used for OvMARK genetic analysis.

| Gene Datasets |  |
| --- | --- |
| GSE26712 | https://www.ncbi.nlm.nih.gov/geo/query/acc.cgi?acc=GSE26712 |
| GSE13876 | https://www.ncbi.nlm.nih.gov/geo/query/acc.cgi?acc=GSE13876 |
| GSE14764 | https://www.ncbi.nlm.nih.gov/geo/query/acc.cgi?acc=GSE14764 |
| GSE30161 | https://www.ncbi.nlm.nih.gov/geo/query/acc.cgi?acc=GSE30161 |
| GSE19161 | https://www.ncbi.nlm.nih.gov/geo/query/acc.cgi?acc=GSE19161 |
| GSE19829 | https://www.ncbi.nlm.nih.gov/geo/query/acc.cgi?acc=GSE19829 |
| GSE26193 | https://www.ncbi.nlm.nih.gov/geo/query/acc.cgi?acc=GSE26193 |
| GSE18520 | https://www.ncbi.nlm.nih.gov/geo/query/acc.cgi?acc=GSE18520 |
| GSE31245 | https://www.ncbi.nlm.nih.gov/geo/query/acc.cgi?acc=GSE31245 |
| GSE9899 | https://www.ncbi.nlm.nih.gov/geo/query/acc.cgi?acc=GSE9899 |
| GSE17260 | https://www.ncbi.nlm.nih.gov/geo/query/acc.cgi?acc=GSE17260 |
| GSE32062 | https://www.ncbi.nlm.nih.gov/geo/query/acc.cgi?acc=GSE32062 |
| TCGA | https://www.cancer.gov/about-nci/organization/ccg/research/structural-genomics/tcga |

**Supplemental table 3:** Differential expression of individual genes in the 10-gene panel correlates with disease-free survival. The OvMark algorithm was used to determine hazard ratios (>1 correlates with poor outcome, <1 correlates with good outcome-blue) and to show statistical significance between high and low expression.

|  | Expression |  |
| --- | --- | --- |
|  | Hazard ratio | p value |
| *ACTB* | 1.115 | 0.0356 |
| *AEBP1* | 1.238 | 4.09E-5 |
| *COL5A1* | 1.229 | 0.00502 |
| *COL11A1* | 1.276 | 9.19E-4 |
| *LOX* | 1.234 | 0.0042 |
| *NECTIN4* | 1.03 | 0.7211 |
| *POSTN* | 1.341 | 1.55E-08 |
| *SNAI1* | 0.9676 | 0.657 |
| *THBS1* | 1.278 | 8.39E-4 |
| *TIMP3* | 1.339 | 1.89E-15 |

**Supplemental table 4:**
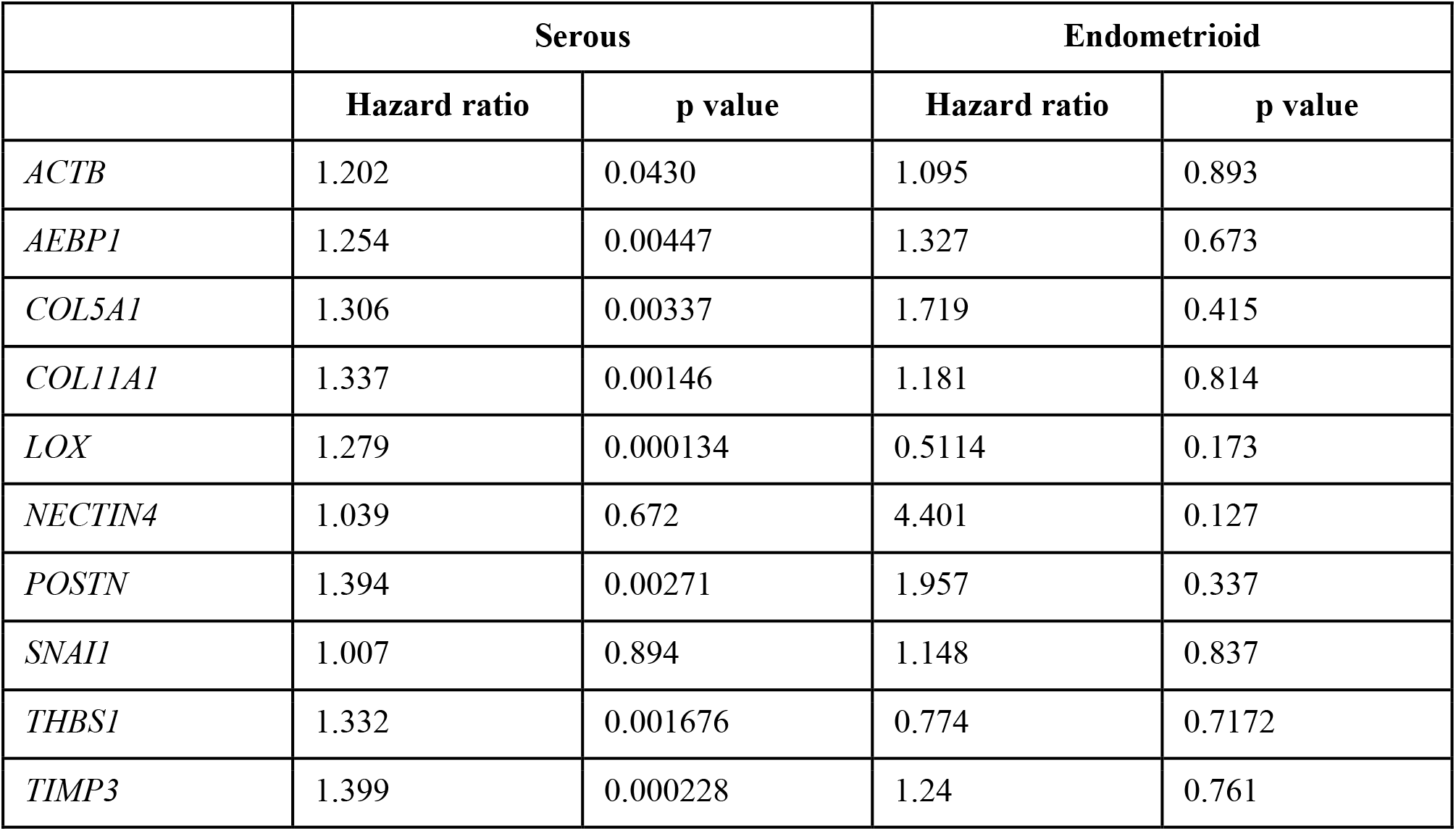
Differential expression of individual genes in the 10-gene panel correlates with disease-free survival in patients with serous ovarian cancer and endometrioid cancer. The OvMark algorithm was used to determine hazard ratios (>1 correlates with poor outcome, <1 correlates with good outcome-blue) and to show statistical significance between high and low expression.

**Supplemental table 5:**
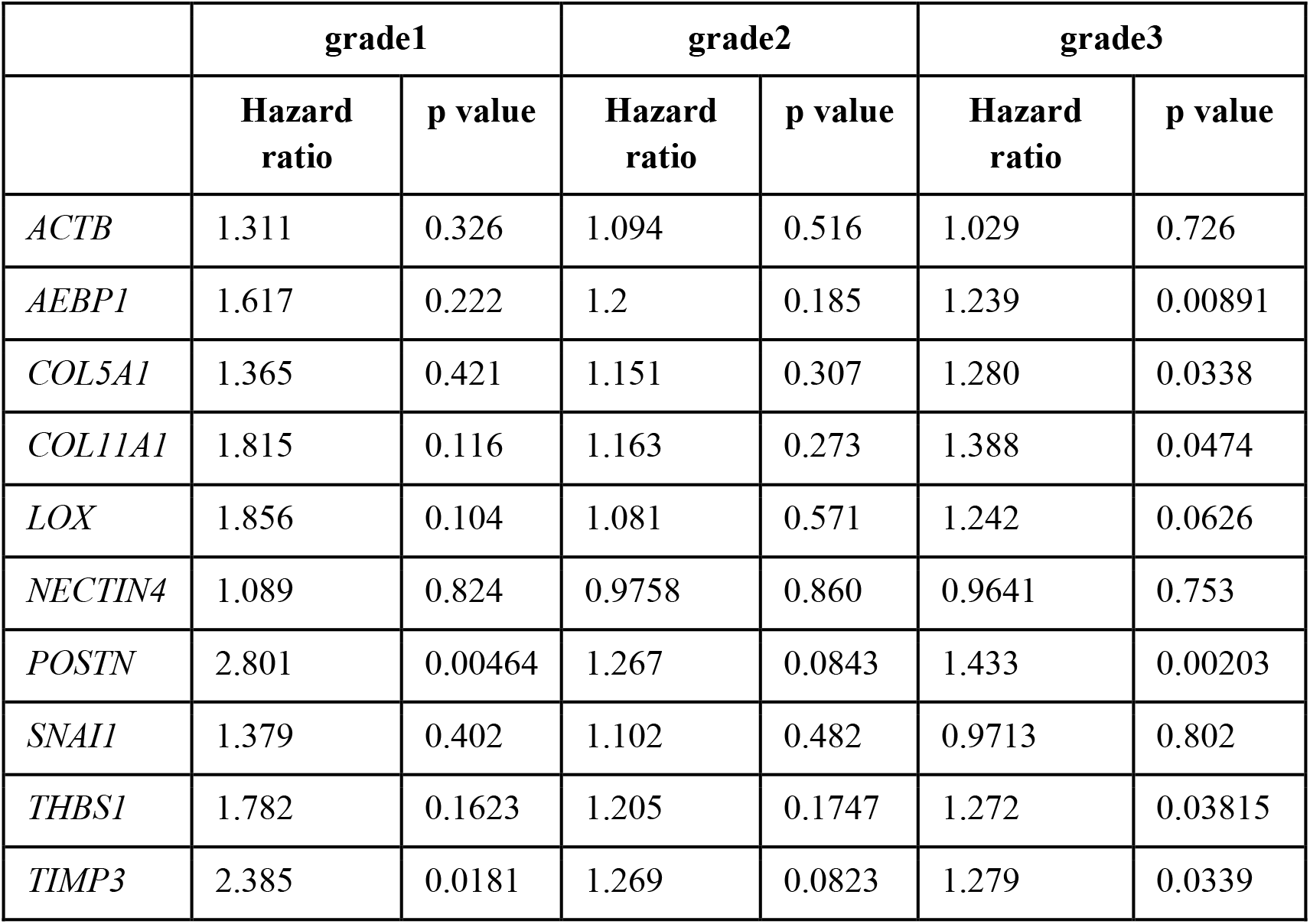
Differential expression of individual genes in the 10-gene panel correlates with disease-free survival in various stages of ovarian cancer development. The OvMark algorithm was used to determine hazard ratios (>1 correlates with poor outcome, <1 correlates with good outcome-blue) and to show statistical significance (red) between high and low expression.

|  | grade1 |  | grade2 |  | grade3 |  |
| --- | --- | --- | --- | --- | --- | --- |
|  | Hazard ratio | p value | Hazard ratio | p value | Hazard ratio | p value |
| *ACTB* | 1.311 | 0.326 | 1.094 | 0.516 | 1.029 | 0.726 |
| *AEBP1* | 1.617 | 0.222 | 1.2 | 0.185 | 1.239 | 0.00891 |
| *COL5A1* | 1.365 | 0.421 | 1.151 | 0.307 | 1.280 | 0.0338 |
| *COL11A1* | 1.815 | 0.116 | 1.163 | 0.273 | 1.388 | 0.0474 |
| *LOX* | 1.856 | 0.104 | 1.081 | 0.571 | 1.242 | 0.0626 |
| *NECTIN4* | 1.089 | 0.824 | 0.9758 | 0.860 | 0.9641 | 0.753 |
| *POSTN* | 2.801 | 0.00464 | 1.267 | 0.0843 | 1.433 | 0.00203 |
| *SNAI1* | 1.379 | 0.402 | 1.102 | 0.482 | 0.9713 | 0.802 |
| *THBS1* | 1.782 | 0.1623 | 1.205 | 0.1747 | 1.272 | 0.03815 |
| *TIMP3* | 2.385 | 0.0181 | 1.269 | 0.0823 | 1.279 | 0.0339 |

**Supplemental table 6:**
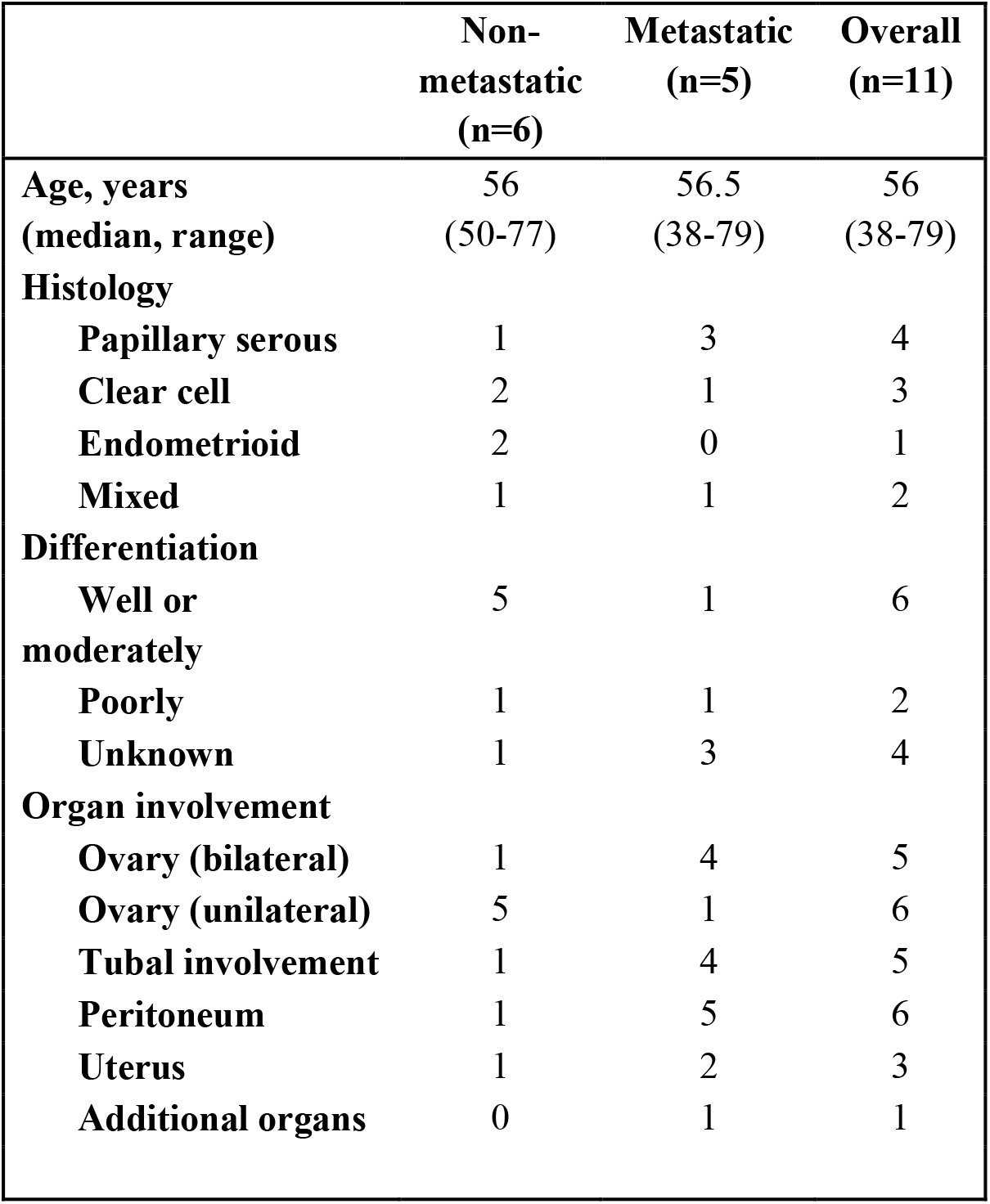
Patient characteristics.

## Abbreviations

*ACTB*: actin-beta
*AEBP1*: AE Binding Protein
ANOVA: analysis of variance
*COL5A1*: collagen type V alpha 1 chain
*COL11A1*: collagen type XI alpha 1 chain
ECM: extracellular matrix
EMT: epithelial-mesenchymal transition
EV: extracellular vesicle
FDA: United States Food and Drug Administration
*LOX*: lysyl oxidase
*NECTIN4*: nectin cell adhesion molecule 4
NTA: nanoparticle tracking analysis
*POSTN*: periostin
PBS: phosphate-buffered saline
RFP: red fluorescence protein
RT-PCR: real-time polymerase chain reaction
sEV: small extracellular vesicle
*SNAI1*: snail family transcriptional repressor 1
TCGA: the Cancer Genome Atlas
*THBS1*: thrombospondin 1
*TIMP3*: tissue inhibitor of metalloproteinase 3
TME: tumor microenvironment
TNM: tumor-node-metastasis staging

